# Maintenance of community function through compensation breaks down over time in a desert rodent community

**DOI:** 10.1101/2021.10.01.462799

**Authors:** Renata M. Diaz, S. K. Morgan Ernest

**Author notes:** **Original submission:** This submission analyzes long-term data on rodent community abundance and energy use from the Portal Project. Sections of this timeseries have been analyzed in numerous other publications, but this is the first to analyze data from 2007-2020 on compensation on experimental and control plots. **Animal welfare:** Rodent censuses were conducted with IACUC approval, most recently under protocol 201808839_01 at the University of Florida. **Open research:** All data and code to reproduce these analyses are archived on Zenodo at https://doi.org/10.5281/zenodo.5544362 and https://doi.org/10.5281/zenodo.5539881. **Analytic methods:** All analyses were conducted in R version 4.0.3.

## Abstract

Understanding the ecological processes that maintain community function in systems experiencing species loss, and how these processes change over time, is key to understanding the relationship between community structure and function and predicting how communities may respond to perturbations in the Anthropocene. Using a 30-year experiment on desert rodents, we show that the impact of species loss on community-level energy use has changed repeatedly and dramatically over time, due to 1) the addition of new species to the community, and 2) a reduction in functional redundancy among the same set of species. Although strong compensation, initially driven by the dispersal of functionally redundant species to the local community, occurred in this system from 1997-2010, since 2010, compensation has broken down due to decreasing functional overlap within the same set of species. Simultaneously, long-term changes in sitewide community composition due to niche complementarity have decoupled the dynamics of compensation from the overall impact of species loss on community-level energy use. Shifting, context-dependent compensatory dynamics, such as those demonstrated here, highlight the importance of explicitly long-term, metacommunity, and eco-evolutionary perspectives on the link between species-level fluctuations and community function in a changing world.

## Introduction

Determining the extent to which community-level properties are affected by species loss, and how and why this changes over time, is key for understanding how communities are structured and how community function may respond to future perturbations (Gonzalez and Loreau 2009). When species are lost from a community, their contributions to community function (e.g. total productivity or resource use) are also directly lost. Community function may be maintained, however, if in the new community context, species that remain perform similar functions to the species that were lost, and compensate for the decline in function directly caused by species loss - i.e., functional redundancy (Walker 1992, 1995; Ernest and Brown 2001; Rosenfeld 2002; Gonzalez and Loreau 2009). When compensation via functional redundancy occurs among consumers with a common resource base, it is consistent with a zero-sum competitive dynamic, in which resources not used by one species are readily absorbed by competitors, and any increases in the abundance of one species must come at the direct expense of others (Van Valen 1973; Ernest et al. 2008).

Because the response of system-level function to species loss is partially determined by degree of functional redundancy in a community, processes that cause functional redundancy to change over time can have important consequences for the long-term maintenance of ecosystem function. Colonization events may buffer community function against species loss, if a community gains species that perform similar functions to the species that were lost (Ernest and Brown 2001; Leibold et al. 2017). The ability of colonization to supply functionally redundant species depends on the species (and traits) present in the broader metacommunity, and on the rate of dispersal supplying appropriate species to local communities (Leibold et al. 2017).

Even without the addition of new species and traits, however, functional redundancy within a consistent set of coexisting species may fluctuate over time. While, in theory, functional redundancy may occur via the special case of complete niche neutrality (e.g. Hubbell 2001), it may also occur in niche-structured systems that contain species that share some traits but differ along other niche axes (Thibault et al. 2010). In these systems, if similar, but non-identical, species respond to environmental change in similar ways, functional overlap can be maintained or even strengthened. However, if niche differences cause species to respond differently to changing conditions, the degree of functional overlap between those species may decline, resulting in a breakdown in compensation (Loreau 2004; Fetzer et al. 2015). Over time, as metacommunity dynamics and changing environmental conditions modulate functional redundancy within a community, the extent to which community function is robust to species loss - and the strength of zero-sum competition - may also be dynamic and context-dependent.

Despite logical conceptual support, and evidence from experimental microcosms (Fetzer et al. 2015), there is little empirical documentation of how, and through which mechanisms, temporal changes in functional redundancy modulate the effect of species loss on ecosystem function in natural assemblages. Although relatively plentiful, observational data cannot unambiguously detect compensation through functional redundancy, and even short-term experiments may not be sufficient to capture temporal variation in compensation (Ernest and Brown 2001; Houlahan et al. 2007). In contrast, long-term manipulative experiments are uniquely suited to address this question. In long-term experiments in which key species are removed from a community over an extended period of time, the impact of species loss on community function can be directly quantified by comparing community function between complete and manipulated assemblages. As metacommunity dynamics and environmental conditions shift over time, long-term monitoring can reveal how these processes contribute to changes in functional redundancy and ecosystem function across different time periods. Due to the financial and logistical resources required to maintain and monitor whole-community manipulations over long timescales, these experiments are rare in natural systems representative of realistic evolutionary, geographic, and environmental constraints (Hughes et al. 2017).

Here, we use a 30-year experiment on desert rodents to investigate how shifts in functional redundancy alter the effect of species loss on community function over time. In this study, kangaroo rats (*Dipodomys* spp.), the largest and competitively dominant species in the rodent community, have been removed from a subset of experimental plots to explore how the loss of key species affects community function, measured as community-level metabolic flux (“total energy use”, or *Etot*) or total biomass (Ernest et al. 2019). For systems of consumers with a shared resource base, such as this community of granivorous rodents, *Etot* reflects the total amount of resources being processed by an assemblage, and total biomass directly reflects standing biomass. Both are important metrics of community function (Lawton 1994; Ernest and Brown 2001). Long-term monitoring of this experiment has documented repeated shifts in the habitat and species composition of this system, resulting in distinct time periods characterized by different habitat conditions and configurations of the rodent community (Christensen et al. 2018). Abrupt reorganization events in community composition occurred in 1997 and in 2010, associated with the establishment and subsequent decline of the pocket mouse *Chaetodipus baileyi. C. baileyi* is similar in size, and presumably other traits, to kangaroo rats, and its establishment in 1996-97 drove a pronounced increase in compensation due to functional redundancy between *C. baileyi* and kangaroo rats (Ernest and Brown 2001; Thibault et al. 2010). Over the course of this experiment, shifting environmental conditions have caused the habitat at the study site to transition from desert grassland to scrub, driving a shift in baseline rodent community composition away from kangaroo rats and favoring other, smaller, granivores (Brown et al. 1997; Ernest et al. 2008). By making comparisons across these time periods, we explored how shifts in community composition and functional overlap among the same species have contributed to long-term changes in the effect of species loss on community function.

## Methods

### The Portal Project

The Portal Project consists of a set of 24 fenced experimental plots located approximately 7 miles east of Portal, AZ, USA, on unceded land of the Chiricahua Apache. Beginning in 1977, kangaroo rats (*Dipodomys spectabilis, D. merriami*, and *D. ordii*) have been experimentally excluded from a subset of these plots (exclosures), while all other rodents are allowed access through small holes cut in the plot fencing. Control plots, with larger holes, are accessible to all rodents, including kangaroo rats. Rodents on all plots are censused via monthly bouts of live-trapping.Each individual captured is identified to species and weighed. For additional details on the site and methodology of the Portal Project, see Ernest et al. (2019).

### Data

We used data for control and exclosure plots from February 1988 until January 2020. The experimental treatments for some plots have changed over time, and we used the subset of plots that have had the same treatments for the longest period of time (Ernest et al. 2019). Four control plots, and five exclosure plots, met these criteria. In order to achieve a balanced sample, we randomly selected four exclosure plots for analysis. We divided the timeseries into three time periods defined by major transitions in the rodent community surrounding the establishment and decline of *C. baileyi* (Ernest and Brown 2001; Christensen et al. 2018). The first time period (February 1988-June 1997) precedes *C. baileyi*’s establishment at site. We defined *C. baileyi*’s establishment date as the first census period in which *C. baileyi* was captured on all exclosure plots (following Bledsoe and Ernest, 2019). During the second time period (July 1997-January 2010), *C. baileyi* was abundant on both exclosure and control plots. This time period ended with a reorganization event in which *C. balieyi* became scarce sitewide. We used January 2010, the midpoint of the 95% credible interval for the date of this reorganization event as estimated in Christensen et al. (2018), as the end date for this time period. The last time period spans from Feburary 2010-January 2020. For each individual rodent captured, we estimated the individual-level metabolic rate using the scaling relationship between individual body mass and metabolic rate *b =* 5.69 * (*m*^0.75^), where *m* is body mass in grams and *b* is metabolic rate (for details, see White et al. 2004). We calculated treatment and species-level energy use as the sum of the appropriate individuals’ metabolic rates, and total biomass as the sum of individuals’ body mass measurements.

### Statistical analysis of rodent community energy use and biomass

Here, we describe analyses for energy use. For biomass, we repeated these analyses substituting biomass values for energy use throughout. For all variables, we combined data for all plots within a treatment in each monthly census period and calculated treatment-level means. This is necessary to calculate compensation, and we treated other variables in the same way to maintain consistency. A provisional plot-level analysis yielded qualitatively equivalent results (Appendix S1). To measure the overall impact of kangaroo rat removal on *Etot*, we calculated a “total energy ratio” as the ratio of treatment-level *Etot* for kangaroo-rat exclosure plots relative to unmanipulated control plots, i.e. *Etot*_*E*_*/Etot*_*C*_ where *Etot*_*E*_ and *Etot*_*C*_ are total energy use on exclosures and controls, respectively (Thibault et al 2010; Bledsoe and Ernest 2019). This ratio is distinct from compensation, which we defined as the proportion of the energy made available by kangaroo rat removal taken up via compensatory increases in energy use by small granivores (all granivores other than kangaroo rats; *Baiomys taylori, C. baileyi, Chaetodipus hispidus, Chaetodipus intermedius, Chaetodipus penicillatus, Perognathus flavus, Peromyscus eremicus, Peromyscus leucopus, Peromyscus maniculatus, Reithrodontomys fulvescens, Reithrodontomys megalotis*, and *Reithrodontomys montanus*). We calculated this as (*SG*_*E*_ *-SG*_*C*_*)/KR*_*C*_, where *SG*_*E*_ and *SG*_*C*_ are the amount of energy used by small granivores (SG) on exclosure and control plots, respectively, and *KR*_*C*_ is the amount of energy used by kangaroo rats (KR) on control plots (Ernest and Brown 2001). To compare these variables across time periods, we used generalized least squares models (GLS; the R package *nlme*; Pinheiro et al. 2020) of the form (*SG*_*E*_ *– SG*_*C*_*)/KR*_*C*_ ∼ *time period*, for compensation, and *Etot*_*E*_*/Etot*_*C*_ ∼ *time period*, for the total energy ratio. We included a continuous-time autoregressive temporal autocorrelation term to account for temporal autocorrelation between values from monthly census periods within each multi-year time period (for details of model selection, see Appendix S2). To evaluate change in baseline community composition over time, we calculated the proportion of treatment-level energy use accounted for by kangaroo rats on control plots in each census period (*KR*_*C*_*/Etot*_*C*_). Proportional energy use is bounded 0-1 and is therefore not appropriate for GLS, so we compared values across time periods using a binomial generalized linear model (GLM) of the form *KR*_*C*_*/Etot*_*C*_ *∼ time period*. Finally, we calculated the proportional energy use accounted for by *C. baileyi* (CB) on exclosure and control plots in each census period (*CB*_*E*_*/Etot*_*E*_ and *CB*_*C*_*/Etot*_*C*_, respectively). *C. baileyi* was not present at the site prior to 1996, and we restricted the analysis of *C. baileyi* proportional energy use to the second two time periods. We compared *C. baileyi* proportional energy use over time and across treatments using a binomial GLM of the form *CB*_*E*_*/Etot*_*E*_ *∼ time period + treatment*. For all models, we calculated estimated means and 95% confidence or credible intervals for time-period (and, for *C. baileyi*, treatment) level values, and contrasts between time periods (and, for *C. baileyi*, treatments), using the R package *emmeans* (Lenth 2021). Analyses were conducted in R 4.0.3 (R Core Team 2020). Data and code are archived at https://doi.org/10.5281/zenodo.5544362 and https://doi.org/10.5281/zenodo.5539881.

## Results

The impact of kangaroo rat removal on community function has changed repeatedly over time, through a combination of abrupt shifts in compensation associated with *C. baileyi*, and long-term changes in baseline community composition sitewide (Figure 1). These dynamics are qualitatively identical whether function is measured as total energy use (Figure 1; Appendix S2) or total biomass (Appendix S3). The first shift coincided with *C. baileyi*’s establishment in the community beginning in 1996-97 (Figure 1D). *C. baileyi* rapidly became dominant on exclosure plots and dramatically increased compensation (Figure 1B). From 1997-2010, small granivores compensated for an average of 58% of kangaroo rat energy use on control plots (95% interval 48-67%), an increase from an average of 18% from 1988-1997 (95% interval 8-29%; contrast *p* < 0.001; for complete results of all models, see Appendix S2) from 1997-2010. With *C. baileyi*’s addition to the community, the total energy ratio (on exclosures relative to controls; Figure 1A) increased from 30% (20-40%) to 71% (62-79%, contrast *p* < 0.014). In the second shift, beginning around 2010, *C. baileyi*’s abundance sitewide dropped precipitously (Figure 1D). *C. baileyi*’s proportional energy use dropped from an average of 72% (65-80%) to 26% (18-35%, contrast *p* < 0.001) on exclosure plots, and from 11% (6-16%) to essentially 0 on control plots (contrast *p* < 0.001). Other species of small granivore did not make compensatory gains to offset the decline in *C. baileyi* (Figure 1B). As a result, compensation declined from an average of 58% (48-67%) to 28% (17-38%, contrast *p* = 0.002), a level not significantly different from the 18% (8-29%, contrast *p* = .44) observed prior to *C. baileyi*’s establishment at the site. Somewhat paradoxically, while the total energy ratio also dropped following *C. baileyi*’s decline, from an average of 71% (62-79%) from 1997-2010 to 50% (40-60%, contrast *p* = 0.0056) from 2010-2020, it remained higher than its average of 30% (20-40%, contrast *p* = 0.0144) from 1988-1997 (Figure 1A). Over the course of the experiment, community composition shifted sitewide. In later years, kangaroo rats accounted for a lower proportion of baseline *Etot* than they did at the beginning of the study (Figure 1C). From 1988-1997, kangaroo rats accounted for 92% (87-97%) of *Etot* on controls; after 1997, this dropped to an average of approximately 70% (1988-1997 compared to later time periods, both *p* = .0004; 1997-2010 and 2020-2020 not significantly different, *p =* .976). Because the proportion of *Etot* directly lost to kangaroo rat removal was smaller from 2010-2020 than from 1988-1997, the total energy ratio was higher from 2010-2020 than it was from 1988-1997 - even though there was not a detectable difference between the two time periods in the proportion of lost energy being offset through compensation.

**Figure 1.**
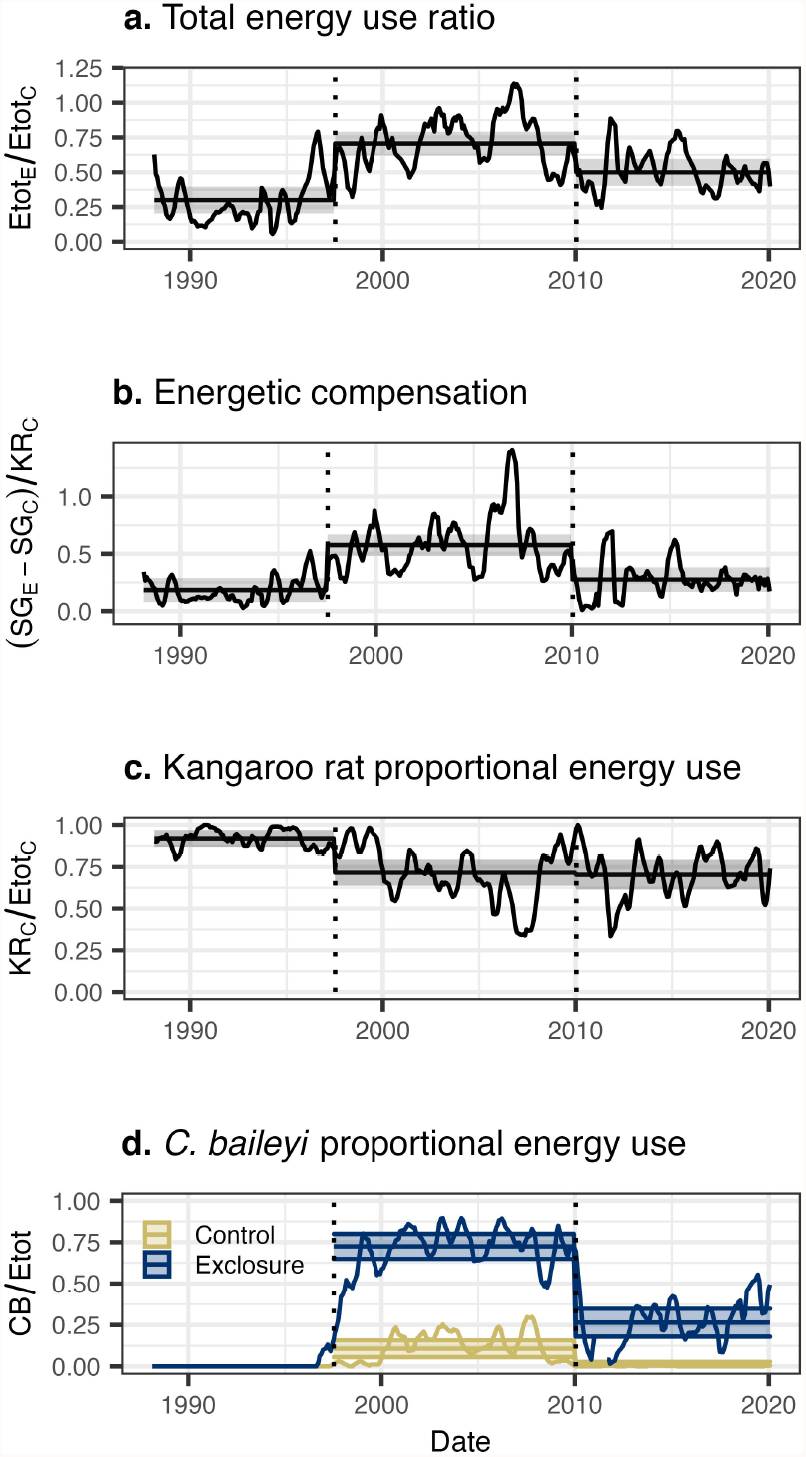
Dynamics of energy use and rodent community composition over time. Lines represent the ratio of total energy use on exclosure plots to control plots (a), 6-month moving averages of energetic compensation (b), and the share of community energy use accounted for by kangaroo rats on control plots (c), and by *C. baileyi* (d), on control (gold) and exclosure (blue) plots. Dotted vertical lines mark the boundaries between time periods used for statistical analysis. Horizontal lines are time-period estimates from generalized least squares (a, b) and generalized linear (c, d) models, and the semitransparent envelopes mark the 95% confidence or credible intervals.

## Discussion

The dynamics of rodent community energy use at Portal illustrate that the role of functional redundancy in buffering community function against species loss fluctuates over time, due to changes in both species composition and in the degree of functional overlap among the same species. The 1997 increase in compensation, driven by *C. baileyi*’s establishment at the site, was a clear and compelling instance of colonization from the regional species pool overcoming limitations on functional redundancy (Ernest and Brown 2001; Leibold et al 2017). Although the small granivore species originally present in the community did not possess the traits necessary to compensate for kangaroo rats, *C. baileyi* supplied those traits and substantially, but incompletely, restored community function. In contrast, following the community reorganization event in 2010, *C. baileyi* remained present in the community, but ceased to operate as a partial functional replacement for kangaroo rats. This is consistent with fluctuating conditions modulating functional redundancy between similar, but non-identical, competitors. Kangaroo rats and *C. baileyi* are relatively similar in size and are demonstrably capable of using similar resources. However, *C. baileyi* prefers different, shrubbier microhabitats than kangaroo rats, and the two groups have been observed to replace each other in adjacent habitats (Ernest and Brown 2001). We suggest that this study site, which has historically been dominated by kangaroo rats, constitutes marginal habitat for *C. baileyi*, and that, while conditions from 1997-2010 aligned sufficiently with *C. baileyi*’s requirements to create appreciable functional redundancy between kangaroo rats and *C. baileyi*, conditions since have caused this redundancy to break down. *C. baileyi*’s decline occurred immediately following a period of low plant productivity and low rodent abundance community-wide, and in the decade following, the site experienced two long and severe droughts (Appendix S4; Christensen et al. 2018). These extreme conditions may themselves have limited *C. baileyi’s* fitness at the site, or the community-wide low abundance event may have temporarily overcome incumbency effects and triggered a community shift tracking longer-term habitat trends (Thibault and Brown 2008; Christensen et al. 2018). Regardless of the proximate cause of *C. baileyi*’s decline, the fact that *C. baileyi* remains in the community, but no longer compensates for kangaroo rats, illustrates that changing conditions can have profound effects on community function by modulating the degree of functional redundancy within a consistent set of species.

While changes in compensation have contributed to changes in community function in this system, changes in compensation alone do not fully account for the long-term changes in the overall impact of kangaroo rat removal on *Etot*. Since 2010, although the ratio of *Etot* on exclosure plots relative to control plots declined coinciding with the breakdown in compensation associated with *C. baileyi*, it remained higher than the levels observed prior to 1997 (Figure 1A). This difference between the first and last time periods cannot be explained by an increase in compensation, as compensation from 2010-2020 was not greater than pre-1997 levels (Figure 1B). Rather, the increase in *Etot* on exclosure plots relative to control plots was the result of a long-term decrease in the contribution of kangaroo rats to *Etot* sitewide. Because kangaroo rats accounted for a smaller proportion of *Etot* on control plots from 2010-2020 than they did prior to 1997, their removal had a smaller impact on community function – even though there was not an increase in the degree to which small granivores compensated for their absence. In fact, the comparable levels of compensation achieved in the decades preceding and following *C. baileyi*’s dominance at the site suggest a relatively stable, and limited, degree of functional overlap between kangaroo rats and the original small granivores (i.e., excluding *C. bailyei*). Niche complementarity, combined with changing habitat conditions, may partially explain this phenomenon. It is well-documented that, while kangaroo rats readily forage in open microhabitats where predation risk can be relatively high, smaller granivores preferentially forage in sheltered microhabitats as an antipredator tactic (Kelt 2011). Over the course of this experiment, the habitat at this study site has transitioned from an arid grassland to a shrubland (Brown et al. 1997). As sheltered microhabitats became more widespread, small granivores may have gained access to a larger proportion of resources and increased their share of *Etot* sitewide. However, kangaroo rats may have continued to use resources in open areas, which would have remained inaccessible to smaller granivores even on exclosure plots. The long-term reduction in the impact of kangaroo rat removal on community function, driven by niche complementarity and consistent niche partitioning, contrasts with the temporary compensatory dynamic driven by functional redundancy with *C. baileyi*. Although changes in the overall effect of species loss are sometimes treated interchangeably with compensation (e.g. Ernest and Brown 2001 compared to Thibault et al. 2010), it is important to recognize that multiple distinct pathways modulate the long-term impacts of species loss on community function. Particularly in strongly niche-structured systems, complementarity effects and fluctuations in functional redundancy may occur simultaneously, with complex and counterintuitive impacts on community function.

Overall, the decadal-scale changes in energy use among the Portal rodents underscore the importance of long-term metacommunity dynamics to the maintenance of community function following species loss (see Leibold et al. 2017). Although a single colonization event may allow for temporary compensation via functional redundancy, as conditions shift, species that once compensated may no longer perform that function (see also Isbell et al. 2011). Particularly if limiting similarity prevents similar competitors from specializing on precisely the same habitats (Rosenfeld 2002), temporary, context-dependent compensation may be common. To maintain compensation over time, multiple colonization events, supplying species that are functionally redundant under different conditions, may be required. Depending on dispersal rates, and the diversity and composition of regional species pools, this may be unlikely or even impossible. At Portal, dispersal limitation introduced a 20-year delay in the compensatory response driven by *C. baileyi*. Theoretically, a new species capable of compensating for kangaroo rats, and better-suited to conditions at the site since 2010, could restore compensation under present conditions – but it is unclear whether this species exists or if it can disperse to this site. As ecosystems globally undergo reductions in habitat connectivity and regional beta diversity, and enter novel climatic spaces, maintenance of community function via functional redundancy may grow increasingly rare and fragile (Dornelas et al. 2014; Williams and Jackson 2007).

Finally, the long-term variability in functional redundancy documented here adds important nuance to our understanding of how zero-sum dynamics operate in natural assemblages. Theories invoking zero-sum dynamics, and tests for compensatory dynamics in empirical data, often treat a zero-sum dynamic as a strong and temporally consistent constraint (Hubbell 2001; Houlahan et al. 2007). In this framing, any resources made available via species loss should immediately be taken up by other species. This is not consistent with the dynamics that occur at Portal, which has seen extended periods of time when resources are available on exclosure plots but are not used. Rather, these results are more consistent with a zero-sum constraint operating at metacommunity or evolutionary scales (Van Valen 1973; Terry and Rowe 2015; Leibold et al. 2017). Over short timescales, or within a closed local assemblage, niche differences may weaken zero-sum effects, especially under fluctuating conditions. However, over larger temporal and spatial scales, dispersal or evolution may supply new species equipped to use available resources - via either functional redundancy, or niche complementarity allowing them to exploit novel niches. A long-term, metacommunity, and even macroevolutionary approach may be necessary to fully understand how zero-sum constraints, functional redundancy, and niche complementarity contribute to the maintenance of community-level function in the face of species extinctions and changing conditions over time.

## Acknowledgements

The Portal Project has been supported by numerous grants from the U.S. National Science Foundation, most recently NSF DEB-1929730. RMD was supported in part by the NSF Graduate Research Fellowship under Grant No. DGE-1315138 and DGE-1842473.

