## Appendix S1 for "Maintenance of community function through compensation breaks down over time in a desert rodent community"

### Appendix S1 - Plot-level analysis

Supplemental information for "Maintenance of community function through compensation breaks down over time in a desert rodent community", by Renata M. Diaz and S. K. Morgan Ernest. In review at Ecology.

Fully annotated code and RMarkdown documents to reproduce these analyses are available at https://doi.org/10.5281/zenodo.5544362 and https://doi.org/10.5281/zenodo.5539881.

### Explanation

In order to calculate energetic compensation and the total energy ratio, we require an estimate for the baseline values of total energy use, kangaroo rat energy use, and small granivore energy use on control plots. Estimating these baselines requires aggregating over between-plot variability among the control plots. For consistency, in the main analysis, we also aggregate across the exclosure plots and focus on treatment-level means throughout. Here, we explore the effect of between-plot variability on our analyses, to the extent possible. We used treatment-level means across control plots to calculate energetic compensation and the total energy ratio, but calculated these quantities separately for each exclosure plot, and conducted analyses including a random effect of plot. We also conducted analyses of *Dipodomys* and *C. baileyi* proportional energy use using plot-level data, again including plot as a random effect. Results were qualitatively the same as using treatment-level means.

### Compensation

#### Model specification and selection

We fit linear mixed-effects models (using the lme function in the R package nlme; Pinheiro et al. 2021) of the form compensation ~ time period with a random effect of plot and temporal autocorrelation structure to account for autocorrelation between monthly census periods within each time period. We compared these to models without the autocorrelation structure, without the random effect, and without the term for time period. The best-fitting model included terms for time period, random effect of plot, and autocorrelation.

**Table S1. Model comparison for compensation.**

| Model.specification | AIC |
|---|---|
| intercept + timeperiod + plot (random effect) + autocorrelation | 1360.207 |
| intercept + timeperiod + plot (random effect) | 1680.916 |
| intercept + timeperiod + autocorrelation | 1409.830 |
| intercept + plot (random effect) + autocorrelation | 1408.362 |
| intercept + plot (random effect) | 1879.126 |
| intercept | 2036.371 |

#### Results

**Table S2. Coefficients from linear mixed-effects model for compensation**

Note that "oera" is the variable name for the term for time period in these analyses.

| | Value | Std.Error | DF | t-value | p-value |
|---|---|---|---|---|---|
| (Intercept) | 0.3451282 | 0.1048354 | 1362 | 3.292096 | 0.0010199 |
| oera.L | 0.0653090 | 0.0373313 | 1362 | 1.749446 | 0.0804392 |
| oera.Q | -0.2845830 | 0.0341063 | 1362 | -8.343990 | 0.0000000 |

**Table S3. Estimates from linear mixed-effects model for compensation**

| Timeperiod | emmean | SE | df | lower.CL | upper.CL |
|---|---|---|---|---|---|
| 1988-1997 | 0.1827673 | 0.1091842 | 3 | -0.1647055 | 0.5302400 |
| 1997-2010 | 0.5774892 | 0.1078860 | 3 | 0.2341478 | 0.9208306 |

| | | | | | |
|---|---|---|---|---|---|
| 2010-2020 | 0.2751282 | 0.1093969 | 3 | -0.0730215 | 0.6232779 |

**Table S4. Contrasts from linear mixed-effects model for compensation**

| Comparison | estimate | SE | df | t.ratio | p.value |
|---|---|---|---|---|---|
| 1988-1997 - 1997-2010 | -0.3947220 | 0.0491845 | 1362 | -8.025330 | 0.0000 |
| 1988-1997 - 2010-2020 | -0.0923609 | 0.0527944 | 1362 | -1.749446 | 0.1873 |
| 1997-2010 - 2010-2020 | 0.3023610 | 0.0496411 | 1362 | 6.090948 | 0.0000 |

### Total energy use

#### Model specification and selection

As for compensation, we fit linear mixed-effects models fitting *total_energy_ratio ~ time period* with a random effect of plot and a temporal autocorrelation term to account for autocorrelation between monthly census periods within each timeperiod. We compared these to models without the autocorrelation term, without the random effect, and without the term for time period. The best-fitting model included terms for time period, random effect of plot, and autocorrelation.

**Table S5. Model comparison for total energy ratio.**

| Model.specification | AIC |
|---|---|
| intercept + timeperiod + plot (random effect) + autocorrelation | 474.8558 |
| intercept + timeperiod + plot (random effect) | 924.1830 |
| intercept + timeperiod + autocorrelation | 507.7842 |
| intercept + plot (random effect) + autocorrelation | 543.5425 |
| intercept + plot (random effect) | 1266.2097 |
| intercept | 1382.7469 |

#### Results

**Table S6. Coefficients from linear mixed-effects model on total energy ratio**

Note that "oera" is the variable name for the term for time period in these analyses.

| | Value | Std.Error | DF | t-value | p-value |
|---|---|---|---|---|---|
| (Intercept) | 0.5018200 | 0.0709701 | 1362 | 7.070865 | 0.0e+00 |
| oera.L | 0.1454309 | 0.0301324 | 1362 | 4.826392 | 1.5e-06 |
| oera.Q | -0.2545852 | 0.0273660 | 1362 | -9.302977 | 0.0e+00 |

**Table S7. Estimates from linear mixed-effects model on total energy ratio**

| Timeperiod | emmean | SE | df | lower.CL | upper.CL |
|---|---|---|---|---|---|
| 1988-1997 | 0.2950508 | 0.0751321 | 3 | 0.0559470 | 0.5341547 |
| 1997-2010 | 0.7096879 | 0.0738511 | 3 | 0.4746606 | 0.9447151 |

| | | | | | |
|---|---|---|---|---|---|
| 2010-2020 | 0.5007212 | 0.0752881 | 3 | 0.2611207 | 0.7403216 |

**Table S8. Contrasts from linear mixed-effects model on total energy ratio**

| Comparison | estimate | SE | df | t.ratio | p.value |
|---|---|---|---|---|---|
| 1988-1997 - 1997-2010 | -0.4146370 | 0.0395736 | 1362 | -10.477622 | 0.0e+00 |
| 1988-1997 - 2010-2020 | -0.2056703 | 0.0426137 | 1362 | -4.826392 | 4.6e-06 |
| 1997-2010 - 2010-2020 | 0.2089667 | 0.0398571 | 1362 | 5.242901 | 5.0e-07 |

### Kangaroo rat proportional energy use

#### Model specification and selection

To compare proportional energy use across time periods, we used binomial generalized linear mixed models (using the glmer function in the R package lme4; Bates et al. 2015), which allowed us to include a random effect of plot.

For *Dipodomys* proportional energy use, we compared models with and without the random effect of plot and with and without a term for timeperiod. The best-fitting model included terms for timeperiod and a random effect of plot.

**Table S9. Model comparison for Dipodomys proportional energy use.**

| Model.specification | AIC |
|---|---|
| intercept + timeperiod + plot (random effect) | 1040.861 |
| intercept + plot (random effect) | 1162.470 |
| intercept + timeperiod | 1108.490 |
| intercept | 1208.081 |

#### Results

**Table S10. Coefficients from GLMER on Dipodomys energy use.**

Note that “oera” is the variable name for the term for time period in these analyses.

| | Estimate | Std. Error | z value | Pr(>\|z\|) |
|---|---|---|---|---|
| (Intercept) | 2.181163 | 0.1305753 | 16.704251 | 0 |
| oera.L | -1.946096 | 0.2664545 | -7.303670 | 0 |
| oera.Q | 1.124620 | 0.1769225 | 6.356572 | 0 |

**Table S11. Estimates from GLMER on Dipodomys energy use.**

Note that estimates are back-transformed onto the response scale, for interpretability.

| Timeperiod | prob | SE | df | asymp.LCL | asymp.UCL |
|---|---|---|---|---|---|
| 1988-1997 | 0.9823009 | 0.0062020 | Inf | 0.9701452 | 0.9944566 |
| 1997-2010 | 0.7795273 | 0.0183934 | Inf | 0.7434769 | 0.8155777 |
| 2010-2020 | 0.7797464 | 0.0208516 | Inf | 0.7388780 | 0.8206149 |

**Table S12. Contrasts from GLMER on Dipodomys energy use.**

Contrasts are performed on the link (logit) scale.

| Comparison | estimate | SE | df | z.ratio | p.value |
|---|---|---|---|---|---|
| 1988-1997 - 1997-2010 | 0.2027736 | 0.0194108 | Inf | 10.4464200 | 0 |
| 1988-1997 - 2010-2020 | 0.2025545 | 0.0217545 | Inf | 9.3109407 | 0 |
| 1997-2010 - 2010-2020 | -0.0002191 | 0.0278048 | Inf | -0.0078811 | 1 |

### C. baileyi proportional energy use

#### Model specification and selection

As for kangaroo rat proportional energy use, we used a binomial generalized linear mixed effects model to compare *C. baileyi* proportional energy use across time periods. Because *C. baileyi* occurs on both control and exclosure plots, we investigated whether the dynamics of *C. baileyi*'s proportional energy use differed between treatment types. We compared models incorporating separate slopes, separate intercepts, or no terms for treatment modulating the change in *C. baileyi* proportional energy use across time periods, i.e. comparing the full set of models:

- *cbaileyi_proportional_energy_use ~ timeperiod + treatment + timeperiod:treatment*
- *cbaileyi_proportional_energy_use ~ timeperiod + treatment*
- *cbaileyi_proportional_energy_use ~ timeperiod*

We also tested a null (intercept-only) model of no change across time periods:

- *cbaileyi_proportional_energy_use ~ 1*

We compared all of these models with and without a random effect of plot.

We found that the best-fitting model incorporated a random effect of plot, and fixed effects for time period and for treatment, but no interaction between them (*cbaileyi_proportional_energy_use ~ timeperiod + treatment*). We therefore proceeded with this model.

**Table S13. Model comparison for C. baileyi proportional energy use.**

| Model.specification | AIC |
|---|---|
| intercept + timeperiod + treatment + timeperiod:treatment + plot (random effect) | 1021.318 |
| intercept + timeperiod + treatment + plot (random effect) | 1020.263 |
| intercept + timeperiod + plot (random effect) | 1042.758 |
| intercept + plot (random effect) | 1321.149 |
| intercept + timeperiod + treatment + timeperiod:treatment | 1166.653 |
| intercept + timeperiod + treatment | 1162.901 |
| intercept + timeperiod | 1869.097 |
| intercept | 2036.489 |

**Results**

**Table S14. Coefficients from GLMER on C. baileyi energy use**

Note that "oera" is the variable name for the term for time period in these analyses, and "oplottype" refers to experimental treatment.

| | Estimate | Std. Error | z value | Pr(>\|z\|) |
|---|---|---|---|---|
| (Intercept) | -2.443643 | 0.2067789 | -11.81766 | 0 |
| oera.L | -1.866286 | 0.1530068 | -12.19740 | 0 |
| oplottype.L | 3.265183 | 0.2913472 | 11.20719 | 0 |

**Table S15. Estimates from GLMER on C. baileyi energy use**

Note that estimates are back-transformed onto the response scale, for interpretability.

| Timeperiod | Treatment | prob | SE | df | asymp.LCL | asymp.UCL |
|---|---|---|---|---|---|---|
| 1997-2010 | Control | 0.0312856 | 0.0116044 | Inf | 0.0085414 | 0.0540297 |
| 1997-2010 | Exclosure | 0.7658194 | 0.0392864 | Inf | 0.6888195 | 0.8428193 |
| 2010-2020 | Control | 0.0023009 | 0.0008486 | Inf | 0.0006378 | 0.0039641 |
| 2010-2020 | Exclosure | 0.1893142 | 0.0364430 | Inf | 0.1178872 | 0.2607412 |

**Table S16. Contrasts from GLMER on C. baileyi energy use.**

Contrasts are performed on the link (logit) scale.

| Comparison | Treatment | estimate | SE | df | z.ratio | p.value |
|---|---|---|---|---|---|---|
| 1997-2010 - 2010-2020 | Control | 2.639326 | 0.2163843 | Inf | 12.1974 | 0 |
| 1997-2010 - 2010-2020 | Exclosure | 2.639326 | 0.2163843 | Inf | 12.1974 | 0 |
