## Appendix S2 for "Maintenance of community function through compensation breaks down over time in a desert rodent community"

### Appendix S2 - Full analytical methods and model results

Supplemental information for "Maintenance of community function through compensation breaks down over time in a desert rodent community", by Renata M. Diaz and S. K. Morgan Ernest. In review at Ecology.

Fully annotated code and RMarkdown documents to reproduce these analyses are available at https://doi.org/10.5281/zenodo.5544362 and https://doi.org/10.5281/zenodo.5539881.

Table of Contents

Compensation 2
- Table S1. Model comparison for compensation. 2
- Table S2. Coefficients from GLS for compensation 2
- Table S3. Estimates from GLS for compensation 2
- Table S4. Contrasts from GLS for compensation 3

Total energy use ratio 4
- Table S5. Model comparison for total energy ratio. 4
- Table S6. Coefficients from GLS on total energy ratio 4
- Table S7. Estimates from GLS on total energy ratio 4
- Table S8. Contrasts from GLS on total energy ratio 4

Kangaroo rat (Dipodomys) proportional energy use 6
- Table S9. Model comparison for Dipodomys proportional energy use. 6
- Table S10. Coefficients from GLM on Dipodomys energy use. 6
- Table S11. Estimates from GLM on Dipodomys energy use. 6
- Table S12. Contrasts from GLM on Dipodomys energy use. 6

C. baileyi proportional energy use 8
- Table S13. Model comparison for C. baileyi proportional energy use. 8
- Table S14. Coefficients from GLM on C. baileyi energy use 8
- Table S15. Estimates from GLM on C. baileyi energy use 9
- Table S16. Contrasts from GLM on C. baileyi energy use. 9

References 10

### Compensation

We fit a generalized least squares (of the form *compensation ~ timeperiod*; note that "timeperiod" is coded as "oera" throughout) using the gls function from the R package nlme (Pinheiro et al. 2021). Because values from monthly censuses within each time period are subject to temporal autocorrelation, we included a continuous autoregressive temporal autocorrelation structure of order 1 (using the CORCAR1 function). We compared this model to models fit without the autocorrelation structure and without the time period term using AIC. The model with both the time period term and the autocorrelation structure was the best-fitting model via AIC, and we used this model to calculate estimates and contrasts using the package emmeans (Lenth 2021).

**Table S1. Model comparison for compensation.**

| Model.specification | AIC |
|---|---|
| intercept + timeperiod + autocorrelation | 69.85023 |
| intercept + autocorrelation | 84.74902 |
| intercept + timeperiod | 157.09726 |
| intercept | 252.74534 |

**Table S2. Coefficients from GLS for compensation**

Note that "oera" is the variable name for the term for time period in these analyses.

| | Value | Std.Error | t-value | p-value |
|---|---|---|---|---|
| (Intercept) | 0.3450313 | 0.0294996 | 11.696141 | 0.0000000 |
| oera.L | 0.0647933 | 0.0524103 | 1.236269 | 0.2172146 |
| oera.Q | -0.2833553 | 0.0477359 | -5.935890 | 0.0000000 |

**Table S3. Estimates from GLS for compensation**

| Timeperiod | emmean | SE | df | lower.CL | upper.CL |
|---|---|---|---|---|---|
| 1988-1997 | 0.1835362 | 0.0520378 | 44.11081 | 0.0786683 | 0.2884041 |
| 1997-2010 | 0.5763899 | 0.0462641 | 47.37851 | 0.4833383 | 0.6694416 |
| 2010-2020 | 0.2751677 | 0.0528010 | 46.75897 | 0.1689314 | 0.3814041 |

**Table S4. Contrasts from GLS for compensation**

| Comparison | estimate | SE | df | t.ratio | p.value |
|---|---|---|---|---|---|
| 1988-1997 - 1997-2010 | -0.3928537 | 0.0689413 | 47.89422 | -5.698378 | 0.0000 |
| 1988-1997 - 2010-2020 | -0.0916315 | 0.0741194 | 45.51740 | -1.236269 | 0.4383 |
| 1997-2010 - 2010-2020 | 0.3012222 | 0.0694989 | 49.52957 | 4.334200 | 0.0002 |

### Total energy use ratio

As for compensation, we fit a generalized least squares of the form *total_energy_ratio ~ timeperiod*, accounting for temporal autocorrelation between monthly censuses within each time period using a continuous autoregressive autocorrelation structure of order 1. We compared this model to models fit without the timeperiod term and/or autocorrelation structure, and found the full (timeperiod plus autocorrelation) model had the best performance via AIC. We used this model for estimates and contrasts.

**Table S5. Model comparison for total energy ratio.**

| Model.specification | AIC |
|---|---|
| intercept + timeperiod + autocorrelation | -132.92138 |
| intercept + autocorrelation | -118.15000 |
| intercept + timeperiod | 13.29396 |
| intercept | 156.85988 |

**Table S6. Coefficients from GLS on total energy ratio**

Note that "oera" is the variable name for the term for time period in these analyses.

| | Value | Std.Error | t-value | p-value |
|---|---|---|---|---|
| (Intercept) | 0.5016731 | 0.0271176 | 18.499880 | 0.0000000 |
| oera.L | 0.1413504 | 0.0477646 | 2.959316 | 0.0033001 |
| oera.Q | -0.2503659 | 0.0429312 | -5.831790 | 0.0000000 |

**Table S7. Estimates from GLS on total energy ratio**

| Timeperiod | emmean | SE | df | lower.CL | upper.CL |
|---|---|---|---|---|---|
| 1988-1997 | 0.2995118 | 0.0475806 | 36.19943 | 0.2030323 | 0.3959913 |
| 1997-2010 | 0.7060960 | 0.0419773 | 38.51943 | 0.6211550 | 0.7910369 |
| 2010-2020 | 0.4994115 | 0.0480066 | 37.62774 | 0.4021956 | 0.5966274 |

**Table S8. Contrasts from GLS on total energy ratio**

| Comparison | estimate | SE | df | t.ratio | p.value |
|---|---|---|---|---|---|
| 1988-1997 - 1997-2010 | -0.4065842 | 0.0623398 | 40.51631 | -6.522060 | 0.0000 |

| | | | | | |
|---|---|---|---|---|---|
| 1988-1997 - 2010-2020 | -0.1998997 | 0.0675493 | 37.12310 | -2.959316 | 0.0144 |
| 1997-2010 - 2010-2020 | 0.2066845 | 0.0626456 | 41.44768 | 3.299267 | 0.0056 |

### Kangaroo rat (Dipodomys) proportional energy use

Proportional energy use is bounded 0-1 and cannot be fit with generalized least squares. We therefore used a binomial generalized linear model of the form *dipodomys_proportional_energy_use ~ timeperiod*. We compared a model fit with a timeperiod term to an intercept-only (null) model using AIC, and found the timeperiod term improved model fit. We used this model for estimates and contrasts.

Note that we were unable to incorporate temporal autocorrelation into generalized linear models, and we prioritized fitting models of the appropirate family over accounting for autocorrelation. Due to the pronounced differences between time periods for these variables, we were comfortable proceeding without explicitly accounting for autocorrelation.

**Table S9. Model comparison for Dipodomys proportional energy use.**

| Model.specification | AIC |
|---|---|
| intercept + timeperiod | 258.3581 |
| intercept | 280.8497 |

**Table S10. Coefficients from GLM on Dipodomys energy use.**

Note that "oera" is the variable name for the term for time period in these analyses. Coefficients are given on the link (logit) scale.

| | Estimate | Std. Error | z value | Pr(>\|z\|) |
|---|---|---|---|---|
| (Intercept) | 1.4032480 | 0.1503201 | 9.335068 | 0.0000000 |
| oera.L | -1.1000833 | 0.2871738 | -3.830723 | 0.0001278 |
| oera.Q | 0.5855493 | 0.2304516 | 2.540878 | 0.0110574 |

**Table S11. Estimates from GLM on Dipodomys energy use.**

Note that estimates are back-transformed onto the response scale, for interpretability.

| Timeperiod | prob | SE | df | asymp.LCL | asymp.UCL |
|---|---|---|---|---|---|
| 1988-1997 | 0.9183528 | 0.0256462 | Inf | 0.8680872 | 0.9686183 |
| 1997-2010 | 0.7160901 | 0.0398537 | Inf | 0.6379782 | 0.7942020 |
| 2010-2020 | 0.7035835 | 0.0456677 | Inf | 0.6140765 | 0.7930905 |

**Table S12. Contrasts from GLM on Dipodomys energy use.**

Contrasts are performed on the link (logit) scale.

| contrast | estimate | SE | df | z.ratio | p.value |
|---|---|---|---|---|---|
| a_pre_pb - b_pre_reorg | 1.4950249 | 0.3942281 | Inf | 3.7922836 | 0.0004 |
| a_pre_pb - c_post_reorg | 1.5557527 | 0.4061251 | Inf | 3.8307227 | 0.0004 |
| b_pre_reorg - c_post_reorg | 0.0607279 | 0.2938992 | Inf | 0.2066282 | 0.9767 |

### C. baileyi proportional energy use

As for kangaroo rat proportional energy use, we used a binomial generalized linear model to compare *C. baileyi* proportional energy use across time periods. Because *C. baileyi* occurs on both control and exclosure plots, we investigated whether the dynamics of *C. baileyi*'s proportional energy use differed between treatment types. We compared models incorporating separate slopes, separate intercepts, or no terms for treatment modulating the change in *C. baileyi* proportional energy use across time periods, i.e. comparing the full set of models:

- *cbaileyi_proportional_energy_use ~ timeperiod + treatment + timeperiod:treatment*
- *cbaileyi_proportional_energy_use ~ timeperiod + treatment*
- *cbaileyi_proportional_energy_use ~ timeperiod*

We also tested a null (intercept-only) model of no change across time periods:

- *cbaileyi_proportional_energy_use ~ 1*

We found that the best-fitting model incorporated effects for time period and for treatment, but no interaction between them (*cbaileyi_proportional_energy_use ~ timeperiod + treatment*). We therefore proceeded with this model.

**Table S13. Model comparison for C. baileyi proportional energy use.**

| Model.specification | AIC |
|---|---|
| intercept + timeperiod + treatment + timeperiod:treatment | 237.7643 |
| intercept + timeperiod + treatment | 231.0963 |
| intercept + timeperiod | 460.8477 |
| intercept | 541.3799 |

**Table S14. Coefficients from GLM on C. baileyi energy use**

Note that "oera" is the variable name for the term for time period in these analyses, and "oplottype" refers to treatment. Coefficients are given on the link (logit) scale.

| | Estimate | Std. Error | z value | Pr(>\|z\|) |
|---|---|---|---|---|
| (Intercept) | -1.574028 | 0.1670168 | -9.424368 | 0 |
| oera.L | -1.409273 | 0.2010398 | -7.009921 | 0 |
| oplottype.L | 2.184896 | 0.2267112 | 9.637355 | 0 |

**Table S15. Estimates from GLM on C. baileyi energy use**

Note that estimates are back-transformed onto the response scale, for interpretability.

| Timeperiod | Treatment | prob | SE | df | asymp.LCL | asymp.UCL |
|---|---|---|---|---|---|---|
| 1997-2010 | Control | 0.1069314 | 0.0258894 | Inf | 0.0561890 | 0.1576737 |
| 1997-2010 | Exclosure | 0.7246076 | 0.0385129 | Inf | 0.6491236 | 0.8000915 |
| 2010-2020 | Control | 0.0160560 | 0.0058224 | Inf | 0.0046444 | 0.0274676 |
| 2010-2020 | Exclosure | 0.2639419 | 0.0428458 | Inf | 0.1799657 | 0.3479181 |

**Table S16. Contrasts from GLM on C. baileyi energy use.**

Contrasts are performed on the link (logit) scale.

| Comparison | Treatment | estimate | SE | df | z.ratio | p.value |
|---|---|---|---|---|---|---|
| 1997-2010 - 2010-2020 | Control | 1.993013 | 0.2843132 | Inf | 7.009921 | 0 |
| 1997-2010 - 2010-2020 | Exclosure | 1.993013 | 0.2843132 | Inf | 7.009921 | 0 |
