## Appendix S3 for "Maintenance of community function through compensation breaks down over time in a desert rodent community"

### Appendix S3 - Biomass analysis

Supplemental information for “Maintenance of community function through compensation breaks down over time in a desert rodent community”, by Renata M. Diaz and S. K. Morgan Ernest. In review at Ecology.

Fully annotated code and RMarkdown documents to reproduce these analyses are available at https://doi.org/10.5281/zenodo.5544362 and https://doi.org/10.5281/zenodo.5539881.

All statistical methods for biomass are identical to the ones for energy use (Appendix S1).

### Compensation

We fit a generalized least squares (of the form *compensation ~ timeperiod*; note that "timeperiod" is coded as "oera" throughout) using the gls function from the R package nlme (Pinheiro et al. 2021). Because values from monthly censuses within each time period are subject to temporal autocorrelation, we included a continuous autoregressive temporal autocorrelation structure of order 1 (using the CORCAR1 function). We compared this model to models fit without the autocorrelation structure and without the time period term using AIC. The model with both the time period term and the autocorrelation structure was the best-fitting model via AIC, and we used this model to calculate estimates and contrasts using the package emmeans (Lenth 2021).

**Table S1. Model comparison for compensation.**

| Model.specification | AIC |
|---|---|
| intercept + timeperiod + autocorrelation | -17.623354 |
| intercept + autocorrelation | -3.297103 |
| intercept + timeperiod | 92.184205 |
| intercept | 207.804481 |

**Table S2. Coefficients from GLS for compensation**

Note that "oera" is the variable name for the term for time period in these analyses.

| | Value | Std.Error | t-value | p-value |
|---|---|---|---|---|
| (Intercept) | 0.3081443 | 0.0290539 | 10.605950 | 0.0000000 |
| oera.L | 0.0711412 | 0.0514131 | 1.383719 | 0.1673549 |
| oera.Q | -0.2799121 | 0.0465252 | -6.016352 | 0.0000000 |

**Table S3. Estimates from GLS for compensation**

| Timeperiod | emmean | SE | df | lower.CL | upper.CL |
|---|---|---|---|---|---|
| 1988-1997 | 0.1435663 | 0.0511419 | 39.28312 | 0.0401458 | 0.2469867 |
| 1997-2010 | 0.5366915 | 0.0452745 | 41.91562 | 0.4453185 | 0.6280646 |
| 2010-2020 | 0.2441751 | 0.0517205 | 41.17937 | 0.1397373 | 0.3486130 |

**Table S4. Contrasts from GLS for compensation**

| Comparison | estimate | SE | df | t.ratio | p.value |
|---|---|---|---|---|---|
| 1988-1997 - 1997-2010 | -0.3931253 | 0.0673811 | 43.22895 | -5.834358 | 0.0000 |
| 1988-1997 - 2010-2020 | -0.1006089 | 0.0727090 | 40.36882 | -1.383719 | 0.3588 |
| 1997-2010 - 2010-2020 | 0.2925164 | 0.0678003 | 44.43055 | 4.314383 | 0.0003 |

### Total biomass ratio

As for compensation, we fit a generalized least squares of the form *total_biomass_ratio ~ timeperiod*, accounting for temporal autocorrelation between monthly censuses within each time period using a continuous autoregressive autocorrelation structure of order 1. We compared this model to models fit without the timeperiod term and/or autocorrelation structure, and found the full (timeperiod plus autocorrelation) model had the best performance via AIC. We used this model for estimates and contrasts.

**Table S5. Model comparison for total biomass ratio.**

| Model.specification | AIC |
|---|---|
| intercept + timeperiod + autocorrelation | -176.57761 |
| intercept + autocorrelation | -162.61339 |
| intercept + timeperiod | -15.98438 |
| intercept | 146.61442 |

**Table S6. Coefficients from GLS on total biomass ratio**

Note that "oera" is the variable name for the term for time period in these analyses.

| | Value | Std.Error | t-value | p-value |
|---|---|---|---|---|
| (Intercept) | 0.4553971 | 0.0272418 | 16.716827 | 0.0000000 |
| oera.L | 0.1454493 | 0.0477989 | 3.042941 | 0.0025257 |
| oera.Q | -0.2531409 | 0.0427343 | -5.923594 | 0.0000000 |

**Table S7. Estimates from GLS on total biomass ratio**

| Timeperiod | emmean | SE | df | lower.CL | upper.CL |
|---|---|---|---|---|---|
| 1988-1997 | 0.2492046 | 0.0476584 | 33.82432 | 0.1523326 | 0.3460765 |
| 1997-2010 | 0.6620857 | 0.0419515 | 35.98516 | 0.5770030 | 0.7471684 |
| 2010-2020 | 0.4549009 | 0.0480215 | 34.98703 | 0.3574107 | 0.5523911 |

**Table S8. Contrasts from GLS on total biomass ratio**

| Comparison | estimate | SE | df | t.ratio | p.value |
|---|---|---|---|---|---|
| 1988-1997 - 1997-2010 | -0.4128811 | 0.0621739 | 38.42746 | -6.640747 | 0.0000 |

| | | | | | |
|---|---|---|---|---|---|
| 1988-1997 - 2010-2020 | -0.2056963 | 0.0675979 | 34.67694 | -3.042941 | 0.0121 |
| 1997-2010 - 2010-2020 | 0.2071848 | 0.0624325 | 39.20390 | 3.318542 | 0.0054 |

#### Kangaroo rat (Dipodomys) proportional biomass

Proportional biomass is bounded 0-1 and cannot be fit with generalized least squares. We therefore used a binomial generalized linear model with no temporal autocorrelation term, of the form *dipodomys_proportional_biomass ~ timeperiod*. We compared a model fit with a timeperiod term to an intercept-only (null) model using AIC, and found the timeperiod term improved model fit. We used this model for estimates and contrasts.

**Table S9. Model comparison for Dipodomys proportional biomass.**

| Model.specification | AIC |
|---|---|
| intercept + timeperiod | 215.2069 |
| intercept | 227.9608 |

**Table S10. Coefficients from GLM on Dipodomys biomass.**

Note that "oera" is the variable name for the term for time period in these analyses. Coefficients are given on the link (logit) scale.

| | Estimate | Std. Error | z value | Pr(>\|z\|) |
|---|---|---|---|---|
| (Intercept) | 1.6149566 | 0.1644937 | 9.817741 | 0.0000000 |
| oera.L | -1.1672395 | 0.3180813 | -3.669626 | 0.0002429 |
| oera.Q | 0.6619048 | 0.2473324 | 2.676175 | 0.0074468 |

**Table S11. Estimates from GLM on Dipodomys biomass.**

Note that estimates are back-transformed onto the response scale, for interpretability.

| Timeperiod | prob | SE | df | asymp.LCL | asymp.UCL |
|---|---|---|---|---|---|
| 1988-1997 | 0.9376458 | 0.0226460 | Inf | 0.8932605 | 0.9820310 |
| 1997-2010 | 0.7454543 | 0.0385025 | Inf | 0.6699909 | 0.8209177 |
| 2010-2020 | 0.7426552 | 0.0437171 | Inf | 0.6569713 | 0.8283392 |

**Table S12. Contrasts from GLM on Dipodomys biomass.**

Contrasts are performed on the link (logit) scale.

| contrast | estimate | SE | df | z.ratio | p.value |
|---|---|---|---|---|---|

| | | | | | |
|---|---|---|---|---|---|
| a_pre_pb - b_pre_reorg | 1.6360275 | 0.4372643 | Inf | 3.741508 | 0.0005 |
| a_pre_pb - c_post_reorg | 1.6507259 | 0.4498349 | Inf | 3.669626 | 0.0007 |
| b_pre_reorg - c_post_reorg | 0.0146984 | 0.3057707 | Inf | 0.048070 | 0.9987 |

### C. baileyi proportional biomass

#### Model specification and selection

As for kangaroo rat proportional biomass, we used a binomial generalized linear model to compare *C. baileyi* proportional biomass across time periods. Because *C. baileyi* occurs on both control and exclosure plots, we investigated whether the dynamics of *C. baileyi*'s proportional biomass differed between treatment types. We compared models incorporating separate slopes, separate intercepts, or no terms for treatment modulating the change in *C. baileyi* proportional biomass across time periods, i.e. comparing the full set of models:

- *cbaileyi_proportional_biomass ~ timeperiod + treatment + timeperiod:treatment*
- *cbaileyi_proportional_biomass ~ timeperiod + treatment*
- *cbaileyi_proportional_biomass ~ timeperiod*

We also tested a null (intercept-only) model of no change across time periods:

- *cbaileyi_proportional_biomass ~ 1*

We found that the best-fitting model incorporated effects for time period and for treatment, but no interaction between them (*cbaileyi_proportional_biomass ~ timeperiod + treatment*). We therefore proceeded with this model.

**Table S13. Model comparison for C. baileyi proportional biomass.**

| Model.specification | AIC |
|---|---|
| intercept + timeperiod + treatment + timeperiod:treatment | 237.6847 |
| intercept + timeperiod + treatment | 231.2374 |
| intercept + timeperiod | 466.4937 |
| intercept + treatment | 346.2154 |
| intercept | 543.7811 |

**Table S14. Coefficients from GLM on C. baileyi biomass.**

Note that "oera" is the variable name for the term for time period in these analyses, and "oplottype" refers to treatment. Coefficients are given on the link (logit) scale.

| | Estimate | Std. Error | z value | Pr(>\|z\|) |
|---|---|---|---|---|
| (Intercept) | -1.538798 | 0.1671239 | -9.207525 | 0 |
| oera.L | -1.403286 | 0.2006948 | -6.992140 | 0 |

| | | | | |
|---|---|---|---|---|
| oplottype.L | 2.270657 | 0.2298594 | 9.878462 | 0 |

**Table S15. Estimates from GLM on C. baileyi biomass**

Note that estimates are back-transformed onto the response scale, for interpretability.

| Timeperiod | Treatment | prob | SE | df | asymp.LCL | asymp.UCL |
|---|---|---|---|---|---|---|
| 1997-2010 | Control | 0.1041331 | 0.0255800 | Inf | 0.0539971 | 0.1542691 |
| 1997-2010 | Exclosure | 0.7425132 | 0.0376727 | Inf | 0.6686761 | 0.8163504 |
| 2010-2020 | Control | 0.0157248 | 0.0057341 | Inf | 0.0044861 | 0.0269634 |
| 2010-2020 | Exclosure | 0.2838438 | 0.0439192 | Inf | 0.1977637 | 0.3699240 |

**Table S16. Contrasts from GLM on C. baileyi biomass.**

Contrasts are performed on the link (logit) scale.

| Comparison | Treatment | estimate | SE | df | z.ratio | p.value |
|---|---|---|---|---|---|---|
| 1997-2010 - 2010-2020 | Control | 1.984546 | 0.2838253 | Inf | 6.99214 | 0 |
| 1997-2010 - 2010-2020 | Exclosure | 1.984546 | 0.2838253 | Inf | 6.99214 | 0 |

**Figure S1. Biomass results**

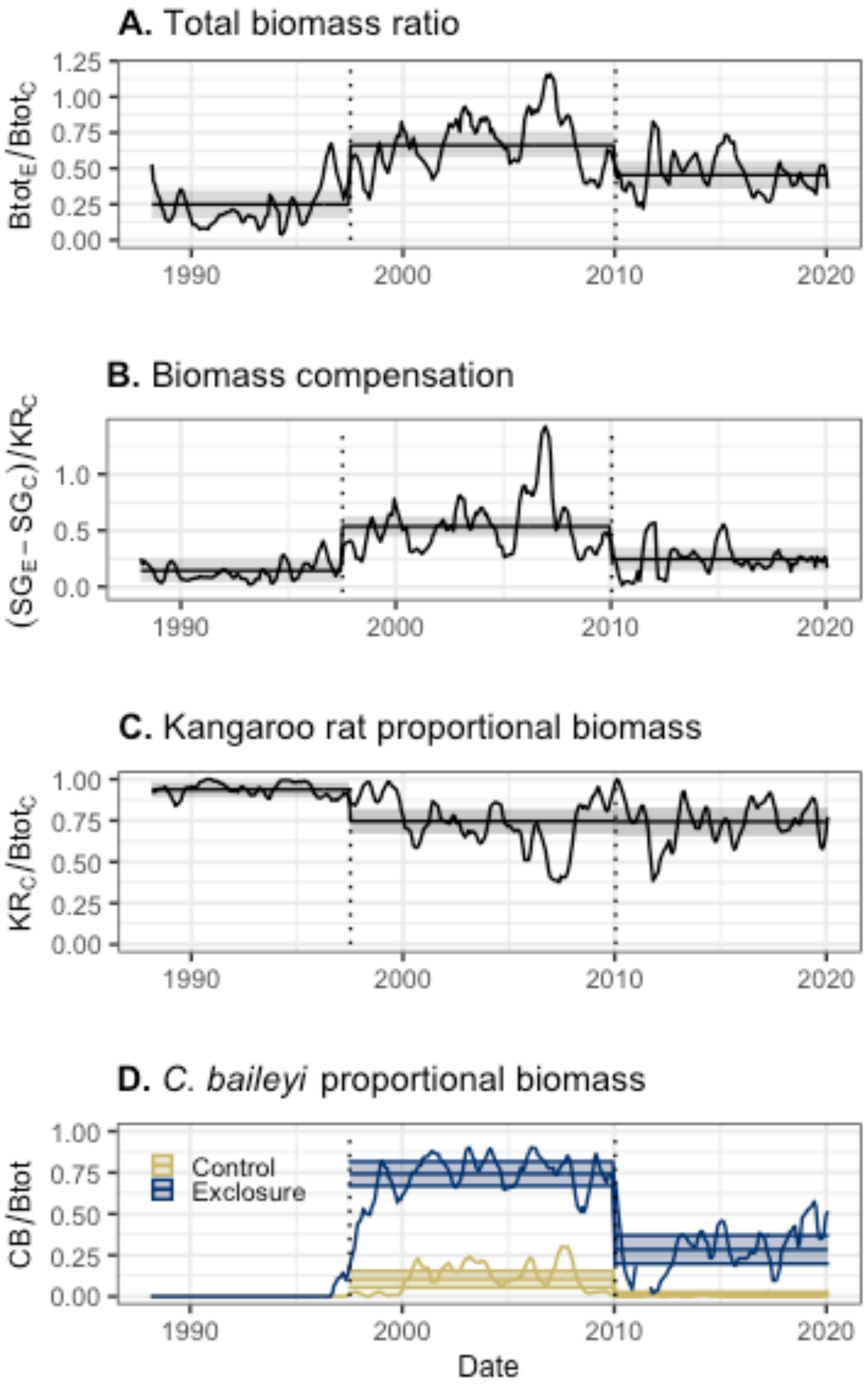

**Figure S1 Legend.**

Dynamics of biomass and rodent community composition over time. Lines represent the ratio of biomass on exclosure plots to control plots (a), 6-month moving averages of biomass compensation (b), and the share of community-wide biomass accounted for by kangaroo rats on control plots (c), and by C. baileyi (d), on control (gold) and exclosure (blue) plots. Dotted vertical lines mark the boundaries between time periods used for statistical analysis. Horizontal lines are time-period estimates from generalized least squares (a, b) and generalized linear (c, d) models, and the semitransparent envelopes mark the 95% confidence or credible intervals.
