## Appendix S4 for "Maintenance of community function through compensation breaks down over time in a desert rodent community"

### Appendix S4 - Covariates of rodent community change

Supplemental information for “Maintenance of community function through compensation breaks down over time in a desert rodent community”, by Renata M. Diaz and S. K. Morgan Ernest. In review at Ecology.

**Appendix S4 Figure S1 - Covariates of rodent community change**

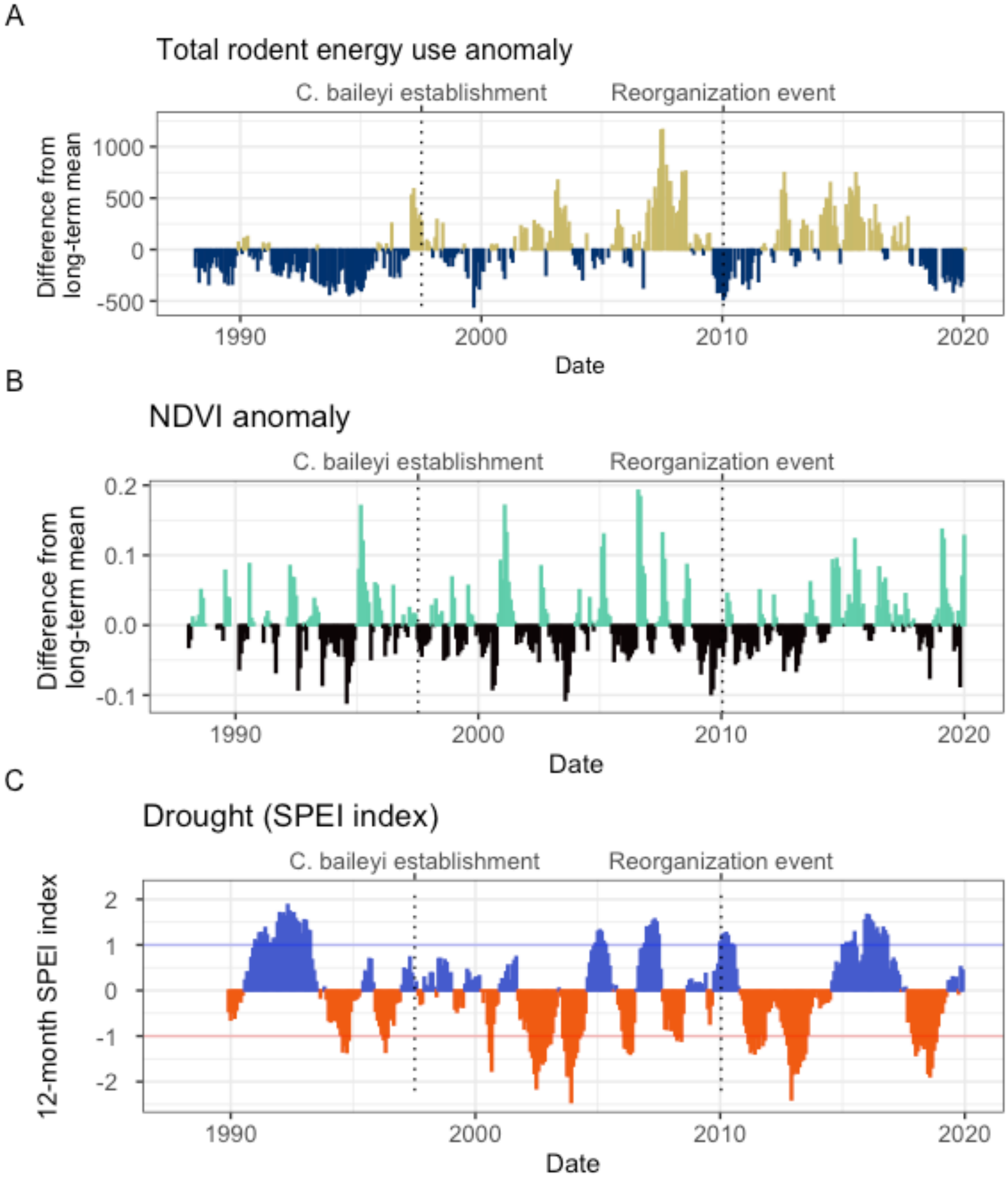

#### Legend

*Figure S1*. Changes in overall community energy use (A), NDVI (B), and local climate (C) surrounding the 2010 shift in rodent community composition. As documented in Christensen et al. (2018), the 2010 transition followed a period of low abundance community-wide (A) and low plant productivity (B). Since 2010, the site has experienced two periods of drought (C) interspersed with an unusually wet period.

Total rodent energy use (A) is calculated as the total energy use of all granviores on control plots ($Etot_C$) in each census period. The anomaly (shown) is calculated as the difference between the total energy use in each census period and the long-term mean of total energy use. Vertical dashed lines mark the dates of major transitions in the rodent community. NDVI anomaly (B) is calculated as the difference between monthly NDVI and the long-term mean for that month. NDVI data were obtained from Landsat 5, 7, and 8 using the ndvi function in the R package portalr (Maesk et al. 2006; Vermote et al. 2016; Christensen et al. 2019). Drought (C) was calculated using a 12-month Standardized Precipitation Evapotranspiraiton index (SPEI) for all months from 1989-2020, using the Thornthwaite method to estimate potential evapotranspiration (using the R package SPEI, Beguería and Vicente-Serrano 2017; Slette et al. 2019; Cárdenas et al. 2021). Values greater than 0 (blue) indicate wetter than average conditions, and values less than 0 (red) indicate drier conditions. Values between -1 and 1 (horizontal lines) are considered within normal variability for a system, while values < -1 constitute drought (Slette et al. 2019).
